# Refining Embedding-Based Binding Predictions by Leveraging AlphaFold2 Structures

**DOI:** 10.1101/2022.08.31.505997

**Authors:** Leopold Endres, Tobias Olenyi, Kyra Erckert, Konstantin Weißenow, Burkhard Rost, Maria Littmann

## Abstract

**Background:** Identifying residues in a protein involved in ligand binding is important for understanding its function. bindEmbed21DL is a Machine Learning method which predicts protein-ligand binding on a per-residue level using embeddings derived from the protein Language Model (pLM) ProtT5. This method relies solely on sequences, making it easily applicable to all proteins. However, highly reliable protein structures are now accessible through the AlphaFold Protein Structure Database or can be predicted using AlphaFold2 and ColabFold, allowing the incorporation of structural information into such sequence-based predictors.

**Results:** Here, we propose bindAdjust which leverages predicted distance maps to adjust the binding probabilities of bindEmbed21DL to subsequently boost performance. bindAdjust raises the recall of bindEmbed21DL from 47±2% to 53±2% at a precision of 50% for small molecule binding. For binding to metal ions and nucleic acids, bindAdjust serves as a filter to identify good predictions focusing on the binding site rather than isolated residues. Further investigation of two examples shows that bindAdjust is in fact able to add binding predictions which are not close in sequence but close in structure, extending the binding residue predictions of bindEmbed21DL to larger binding stretches or binding sites.

**Conclusion:** Due to its simplicity and speed, the algorithm of bindAdjust can easily refine binding predictions also from other tools than bindEmbed21DL and, in fact, could be applied to any protein prediction task.

## Introduction

The correct functioning of proteins is only possible under specific conditions often involving the binding of specific molecules, called *ligands*. As such, the precise identification of residues, involved in ligand binding, is a crucial step in unravelling the function of a protein and serves as a stepping-stone for fields like drug discovery and development [1, 2]. While high-throughput sequencing ensures a constant increase in the availability of experimentally verified sequences, the identification of binding sites is experimentally involved and cannot keep up. As such the field currently relies on predictive methods to bridge this sequence-annotation gap [3].

Traditionally, Machine Learning methods rely on evolutionary information as represented in multiple sequence alignments (MSAs) to predict binding residues [4-6]. Alternatively, homology-based inference transfers information about, e.g., binding residues from sequence-similar proteins with known annotations to uncharacterized proteins [7]. Template-based methods consider structural similarity instead of sequence similarity and therefore require structures with binding annotations [8, 9]. All those approaches are time-consuming and often limited by the small number of available annotated sequences and structures. The emergence of protein Language Models (pLMs) partially addresses this issue. Based on ideas from the field of Natural Language Processing (NLP), pLMs are trained on large unlabeled protein sequence corpora and implicitly learn to encode protein features into a latent vector space [10-12]. Leveraging those vector representations (called *embeddings*) of protein sequences as input to downstream models allows to transfer the encapsulated information to various tasks resulting in high quality predictions for different aspects of protein structure and function (*transfer learning*) [13-19].

One such model is bindEmbed21DL [16], which uses ProtT5 embeddings [10] for each residue to quickly and accurately predict residues binding to small molecules, metal ions, or nucleic acids (DNA and RNA). bindEmbed21DL only requires single sequence input and makes fast predictions outperforming MSA-based predictors [16]. While achieving good performance, bindEmbed21DL does not integrate structural information. Therefore, it is limited to predicting binding residues in the sequence without any information on whether those residues could form a binding site or are isolated in 3D space. While previously inaccessible, the release of AlphaFold2 [20], ColabFold [20] and the AlphaFold Protein Structure Database [21] (AFDB) enables access to high-quality protein structure predictions which can be used to extend sequence-based methods to integrate structural information and subsequently refine predictions.

Here, we propose *bindAdjust*, a simple and fast method leveraging (predicted) protein structures to refine binding predictions of bindEmbed21DL. The refinement algorithm combines the (predicted) distance map and the predicted binding probabilities of a protein to calculate a per-residue *structure bonus*. This bonus is used to adjust the probabilities generated by bindEmbed21DL accordingly. Notably, bindAdjust does not need any additional training apart from hyper-parameter selection and, thus, provides a simple approach to refine binding residue predictions from sequence-based methods.

## Methods

### Data set

The data set was taken from bindEmbed21DL [16]. We removed four sequences due to non-native amino acids in those, which were not considered during structure prediction. Details can be found in Table S1 in the Supporting Online Material (SOM).

### Binding predictions

We computed per-ligand, per-residue binding predictions using bindEmbed21DL [16]. bindEmbed21DL is a convolutional neural network, relying on protein embeddings generated with the Protein Language Model (pLM) ProtT5-XL-UniRef50 [10] (*ProtT5*) as input [16]. bindEmbed21DL provides three output probabilities for each residue in a protein, indicating whether this residue binds to metal ions, nucleic acids, or small molecules. A residue is considered binding to ligand X if the probability p(X)?0.5 [16].

### 3D structures

To obtain three-dimensional (3D) structures for all proteins, we used ColabFold [20], which is based on AlphaFold2 [22]. At the time of writing (June 2022), AlphaFold2 is the state-of-the-art protein structure prediction model. It uses a combination of evolutionary information in the form of multiple sequence alignments (MSAs), templates, and database lookups to predict the folded structure of a protein based on its sequence [22]. ColabFold extends upon vanilla AlphaFold2 through early stopping, faster database lookups, and more comprehensive MSA computations [20]. We derived distance maps from the generated structure files. For subsequent algorithmic input, we used the location of C-Alpha atoms as central amino acid position. ColabFold allows easy generation of high-quality structure predictions on GPUs with 8GB of video RAM. However, with the fourth release of the AlphaFold Protein Structure Database (AFDB) [21] containing 200 million protein structures, this step becomes obsolete for most protein sequences known today.

### bindAdjust

bindAdjust algorithmically adjusts the predicted binding probability of bindEmbed21DL for each ligand class and each protein. To compute the respective adjustment value for one residue, bindAdjust incorporates the distances to all other residues in the protein and their binding probabilities. Distances were extracted from the generated distance maps based on the predicted ColabFold structures. For each residue *i* and each ligand *l* in the sequence and each of the three binding probabilities, we computed a positive *structure bonus* (Eqn. 1) and added it (regularized by a coefficient *C* > 0) to the original bindEmbed21DL binding probability (Eqn. 2). Generally, the computed bonus increases the predicted binding probabilities of residues that are close to residues with high binding probabilities. A visual example is shown in Fig. S1.

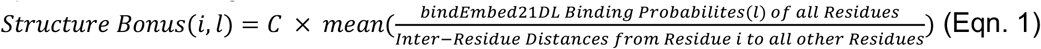

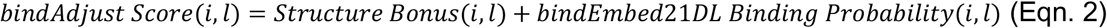

### Runtime

We tested the runtime of bindAdjust on a low performance CPU (2.6 GHz 6-Core Intel Core i7 using 16 Gigabytes of RAM). Calculating the structure bonus values for all three ligand classes of all 1010 proteins and writing them to disk only took around 5 seconds, resulting in a runtime of around 5 milliseconds per protein. We excluded the time for computing the structure and binding predictions from the runtime analysis as with the release of the fourth version of AlphaFold Protein Structure Database (AFDB), 200 million protein structures are available [21] for download and tools like bio-embeddings [23] and LambdaPP [24] allow for fast and easy bindEmbed21DL predictions.

### Bonus coefficient C optimization

The coefficient C was optimized for each ligand class individually using the development set (Table S1). C determines the influence of the structure bonus on the binding probability, thus, substantially influencing the performance of bindAdjust. We tested values for C between 0 and 200 and benchmarked the binarized bindAdjust scores on the development set. Binarized scores were derived by applying a cutoff on the bindAdjust scores (Eqn. 2, more information given below). We used C=38 to evaluate the performance on the test set because it performed well for all ligand classes.

### Label Binarization

bindAdjust produces continuous, per-residue scores (Eqn. 2) based on the predicted bindEmbed21DL output probabilities. To evaluate against the binary binding labels (0: non-binding, 1: binding), we binarized the scores using a cutoff. For each value of C, we identified the cutoff leading to an average precision (Eqn. 3) of 50% of bindAdjust on the development set. All continuous bindAdjust scores were mapped to binary non-binding/binding if the score was smaller/larger than the defined cutoff. We chose a target precision of 50% to be able to compare performances for different values for C and other methods.

### Sequence-Based Refinement

To in ensure that bindAdjust uses information encoded the distance map/3D structure and does not solely rely on information about residue distance in sequence, we implemented a refinement method based on distance in sequence not structure. Analogously to bindAdjust, we calculated a bonus, with the difference that it is based on the position difference of two residues in the sequence (see Table S3 for an example). The distance between two neighboring residues was set to 1. We hypothesize that a sufficiently large performance drop for the sequence-based refinement compared to bindAdjust would indicate that bindAdjust uses additional information beyond the sequence to refine binding residue predictions.

### Performance Metrics and Evaluation

To assess the performance of the refinement achieved by bindAdjust, we classified each output into one of four categories: True Positives (TP) are the number of residues correctly predicted as binding. True Negatives (TN) are residues that have been rightly predicted to be non-binding. False Positives (FP) are the number of residues incorrectly predicted as binding, while False Negatives (FN) are the number of residues wrongly predicted as non-binding. Based on those measures, standard performance metrics could be calculated for each protein, namely precision (Eqn. 3), recall (Eqn. 4), and F1 score (Eqn. 5).

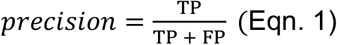

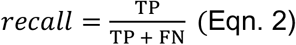

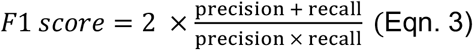

If the model does not predict any residue of a protein as binding (i.e., no bindAdjust score > cutoff), a special case arises where the selected metrics are not defined. In this case, the protein was omitted from the performance evaluation, i.e., performance metrics were calculated for proteins with at least one binding prediction (correct or incorrect). To track the number of proteins included in the evaluation, bindEmbed21DL introduced CovOneBind as the fraction of proteins with at least one residue predicted as binding [16] (Eqn. 6).

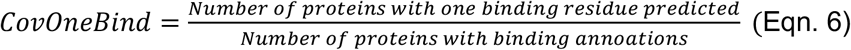

To account for sampling variance, we computed standard errors as the standard deviations of these metrics from 10,000 bootstrapping samples.

### Visualization of binding predictions

We used PyMol [25] to visualize binding predictions of bindAdjust and bindEmbed21DL [16]. On the predicted protein 3D structure of ColabFold [20], we highlighted TP, FP, FN, and TN using the binding annotations of BioLiP [26] as ground truth. This allowed us to visually inspect whether (predicted) binding residues form spatial clusters on the 3D structure [20].

## Results and Discussion

### Recall improves for small molecule binding

603 of the 1010 proteins of the development set have at least one residue annotated to bind to a small molecule in BioLiP [26]. For those proteins, bindEmbed21DL achieves recall=37±1% at a precision of about 50% making predictions for 588 proteins (CovOneBind=90%; Fig. S2, Table S4). bindAdjust improves upon this recall at the same level of precision depending on the choice of C (Table S4, Fig. S3A). Increasing C correlates consistently with an increase in average recall and a respective decrease in CovOneBind. For C>70, bindAdjust cannot achieve the target precision of 50% anymore (Fig. S3A). We choose C=38 and C=66 for further analyses (Fig. S2).

Since bindAdjust achieves an increase in recall by accepting a decrease in CovOneBind, the overall increase could stem from removing all binding predictions for proteins with low recall. To assess whether this is the case, we compare the prediction performance for the 479 proteins, for which both bindEmbed21DL and bindAdjust at C=38 predict at least one residue as small molecule binding. C=38 is chosen because this value leads to an increase in recall with an acceptable trade-off in terms of CovOneBind compared to C=66 (Fig. S2). bindEmbed21DL achieves a mean recall of 40±1%, while bindAdjust achieves a recall of 45±1% at the same level of precision for the same proteins (Fig. S4). This analysis shows that the increase in performance achieved by bindAdjust does not originate from the removal of proteins with subpar prediction performance and, thus, originates from improvement in the prediction quality.

We also benchmark bindAdjust using C=38 and C=66 against bindEmbed21DL on the test set (TestSet220; Fig. 1). Using the cutoff which results in a mean precision of 50% on the development set, bindEmbed21DL achieves recall=38±2% and precision=58±2% making predictions for 202 of the 220 small binding proteins (CovOneBind=92%) while bindAdjust achieves a similar recall and precision but at a lower CovOneBind (Fig. 1). Hence, bindAdjust cannot improve upon the predictions of bindEmbed21DL on the test set. This could be due to the surprisingly higher performance of bindEmbed21DL on its test set than on its validation set [16]. Also, while achieving similar precision, the performance values for bindEmbed21DL and bindAdjust on the test set are not directly comparable when using the cutoffs leading to precision=50% on the development set.

**Fig. 1:**
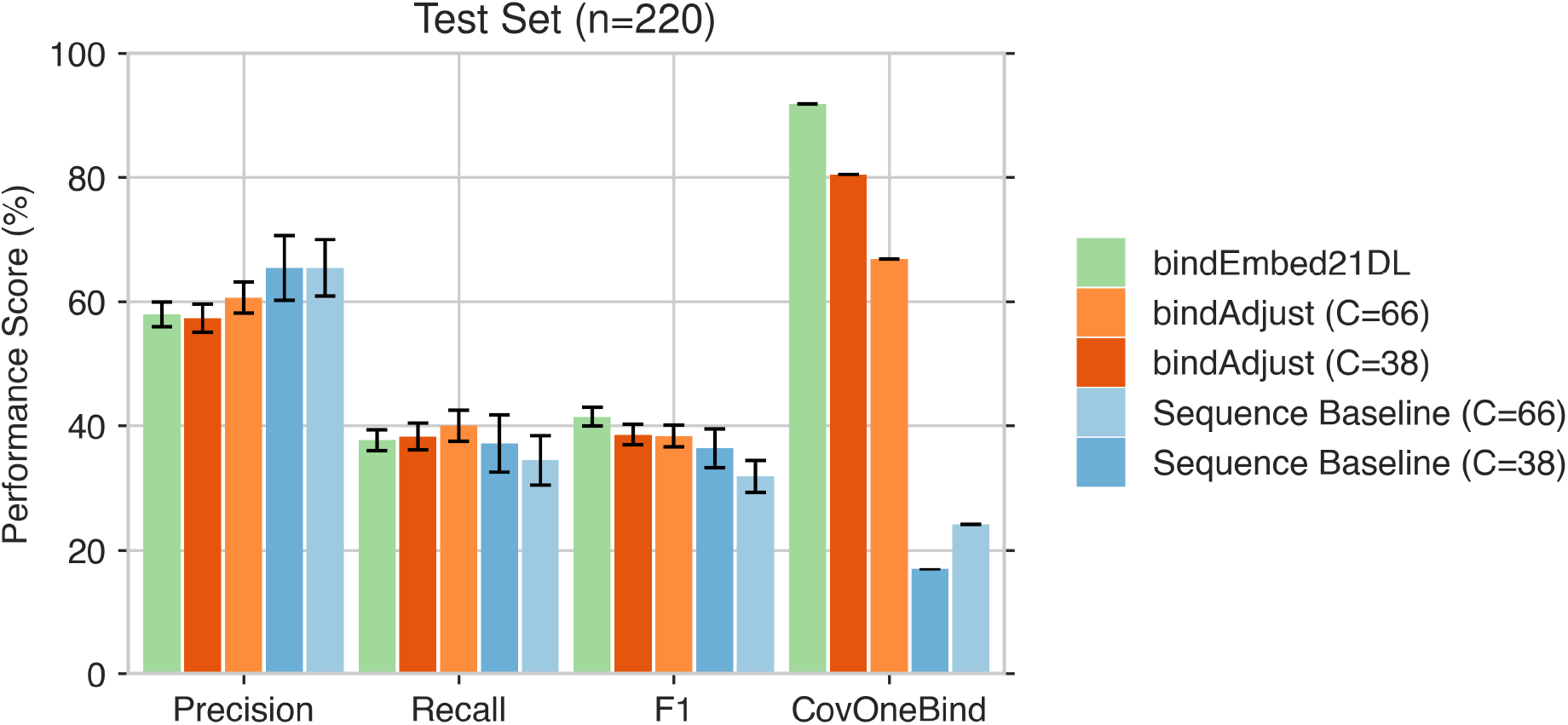
Average performance metrics of bindEmbed21DL, the sequence-based refinement, and bindAdjust for test and parameter settings for the prediction of small molecule binding. *<u>TestSet220</u>*: 220 proteins binding to small lecules from the test set. For the test set, the same cutoffs to classify a residue as (non-)binding were chosen as for the elopment set. Standard errors are given as error estimates. Using the same cutoffs as for the development set, dAdjust and bindEmbed21DL achieve a similar performance in terms of precision, recall, and F1 with bindAdjust making dictions for less proteins than bindEmbed21DL.

If we choose cutoffs for bindEmbed21DL and bindAdjust that lead to a precision of 50% on the test set, we observe that the recall improves by five and six percentage points for C=38 and C=66, respectively, accepting a drop in CovOneBind (Table S4). For the set of 199 proteins (TestSet199) where both bindAdjust and bindEmbed21DL predicted at least one residue as binding, bindAdjust achieves recall=52±2% significantly improving over bindEmbed21DL (p=0.0003, Table S5, Fig. 2). So, while the improvement of bindAdjust is not evident when using the cutoffs optimized on the development set, this seems to be rather due to the test set performance of bindEmbed21DL not reflecting its real performance [16] and does not mean that bindAdjust cannot refine binding residue predictions in general.

**Fig. 2:**
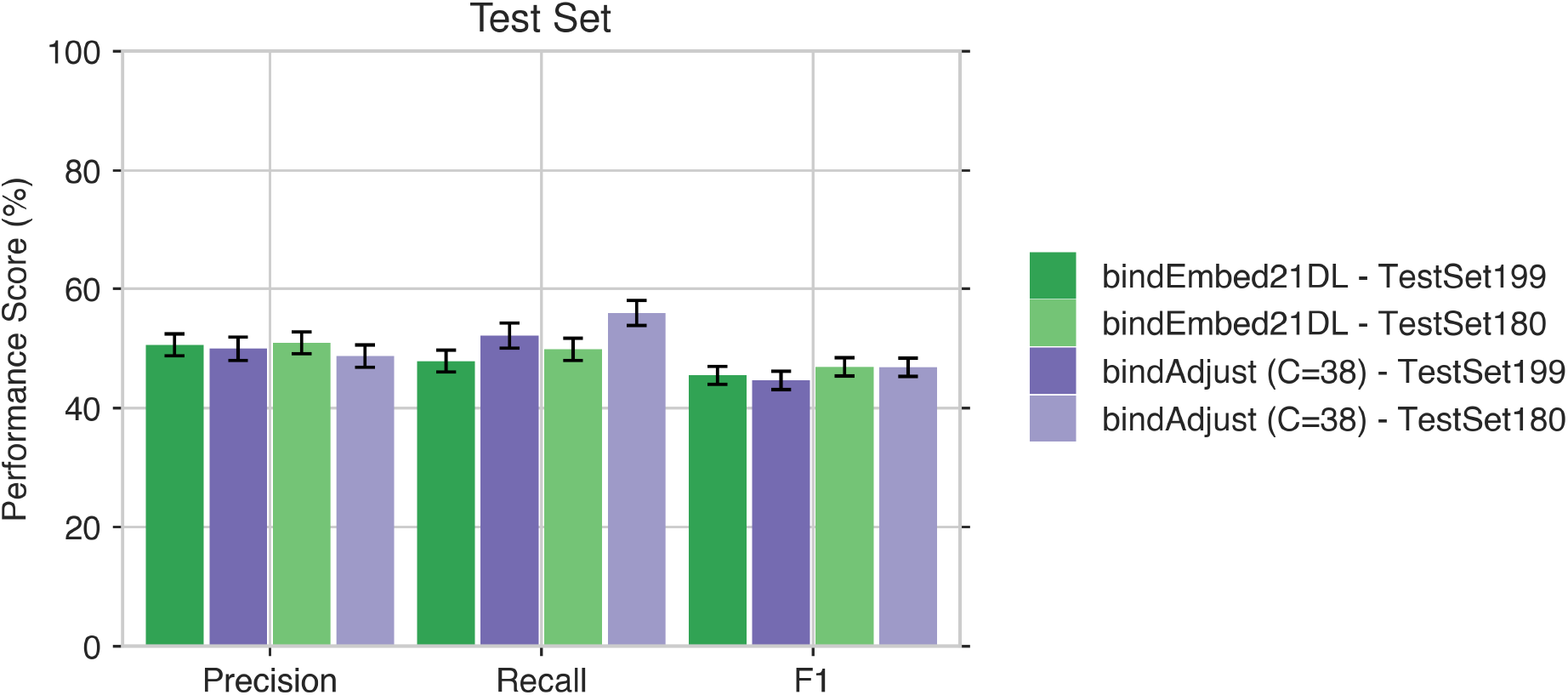
Average performance metrics of bindEmbed21DL and bindAdjust(C=38) for the prediction of small molecule ding on protein subsets identified by bindAdjust. *<u>TestSet199</u>*: 199 proteins of the test set binding to small molecules which both bindEmbed21DL and bindAdjust(C=38) predict at least one residue as small binding. *<u>TestSet180</u>*: 180 teins of the test set binding to small molecules for which bindAdjust(C=66) predicts at least one residue as small binding. the test set, the cutoffs leading to 50% precision were used to classify a residue as (non-) binding. Standard errors are en as error estimates. Using *<u>TestSet188</u>*, both bindAdjust(C=38) and bindEmbed21DL perform better than for*<u>tSet199</u>*. This demonstrates that bindAdjust and large values for C can be used to identify a subset of proteins for which h analyzed models predict small binding exceptionally well.

### bindAdjust identifies set of particularly good predictions for metal ions and nucleic acids

Performing similar analyses for the prediction of binding to metal ions and nucleic acids reveals that bindAdjust can also improve the recall for those ligand classes for C=66 (results for DevSet454 – nucleic acid binding -in Table S6 and for DevSet108 – binding to metal ions – in Table S7). However, focusing on the set of proteins for which both bindEmbed21DL and bindAdjust made a prediction, the recall of bindEmbed21DL either reaches the same level as bindAdjust (for metal ions; Table S6) or even exceeds it (for nucleic acids; Table S7). Therefore, bindAdjust rather allows to identify a subset of proteins with particularly good binding predictions by bindEmbed21DL than improving predictions for individual proteins. While not directly enabling a refinement of predictions, bindAdjust can be used as a filter to identify good predictions of bindEmbed21DL especially in terms of recall. Since the refinement of bindAdjust is based on structural information, we assume that the set of good predictions, identified by bindAdjust form spatial patterns indicating the prediction of an actual binding site rather than isolated residues. This could also explain why bindAdjust cannot improve those predictions: bindEmbed21DL correctly identifies the complete binding site and therefore bindAdjust cannot identify any additional binding residues in this site. More details can be found in the SOM Section 2.3.

### bindAdjust leverages spatial information not available from sequence

The sequence-based refinement method only uses distance in sequence not in structure to refine predictions. For C=38, the recall for predicting binding to small molecules improves over bindEmbed21DL by four percentage points, but the sequence-based refinement cannot reach the same level of performance as bindAdjust (Table S4). At C=66, the sequence-based method does not reach the target precision level of 50% (Table S4). For both binding to metal ions and binding to nucleic acids, the sequence-based refinement cannot improve the recall of bindEmbed21DL (Table S6 for metal ions, Table S7 for nucleic acids). Using information about residues close in sequence space can apparently help in refining the predictions to a certain extent. However, incorporating structure information as done by bindAdjust is needed to obtain a large improvement. In addition, the incorporation of structural information allows for the identification of binding sites (i.e., groups of binding residues close in space) not possible from sequence alone. Hence, bindAdjust incorporates new information and allows for a performance boost compared to only using protein sequences.

### Case study: bindAdjust leverages spatial relations between binding residues

From the development set, we visualized two small binding proteins with an increase in F1 score compared to bindEmbed21DL showcasing how bindAdjust can improve the binding predictions from bindEmbed21DL by taking residues close in 3D space into account. For bindEmbed21DL, a cutoff of 0.65 was applied to classify a residue as binding. For the sequence-based refinement and bindAdjust, C was set to 38 and cutoffs of 1.35 and 1.15 were applied, respectively. We chose these cutoffs as they corresponded to an average precision of 50% on the development set (Table S4).

The rhodothermus marinus cytochrome c (Fig. 3 first row; UniProt ID B3FQS5 [27]) is annotated to have 28 residues binding to a small molecule according to BioLiP [26]. bindEmbed21DL predicts 16 residues as binding of which 14 are correct (Recall=50%, Precision=88%), i.e., bindEmbed21DL predicts too few residues but at a high precision (Fig. 3A). Applying the sequence-based refinement method, the recall improves to 61% by adding five new binding predictions (Fig. 3B) of which three are correct (Precision=81%). bindAdjust adds seven residues to the predictions of bindEmbed21DL resulting in 23 binding residues of which 20 were correct, i.e., increasing the recall to 71% at a similar level of precision (Precision=87%; Fig. 3C). Mapping those predictions to the predicted 3D structure clearly shows that the newly added residues are in fact close to the original predictions from bindEmbed21DL (Fig. 3C), while not all being close in sequence space (otherwise, they would have been identified with the sequence-based method; Fig. 3B).

**Fig. 3:**
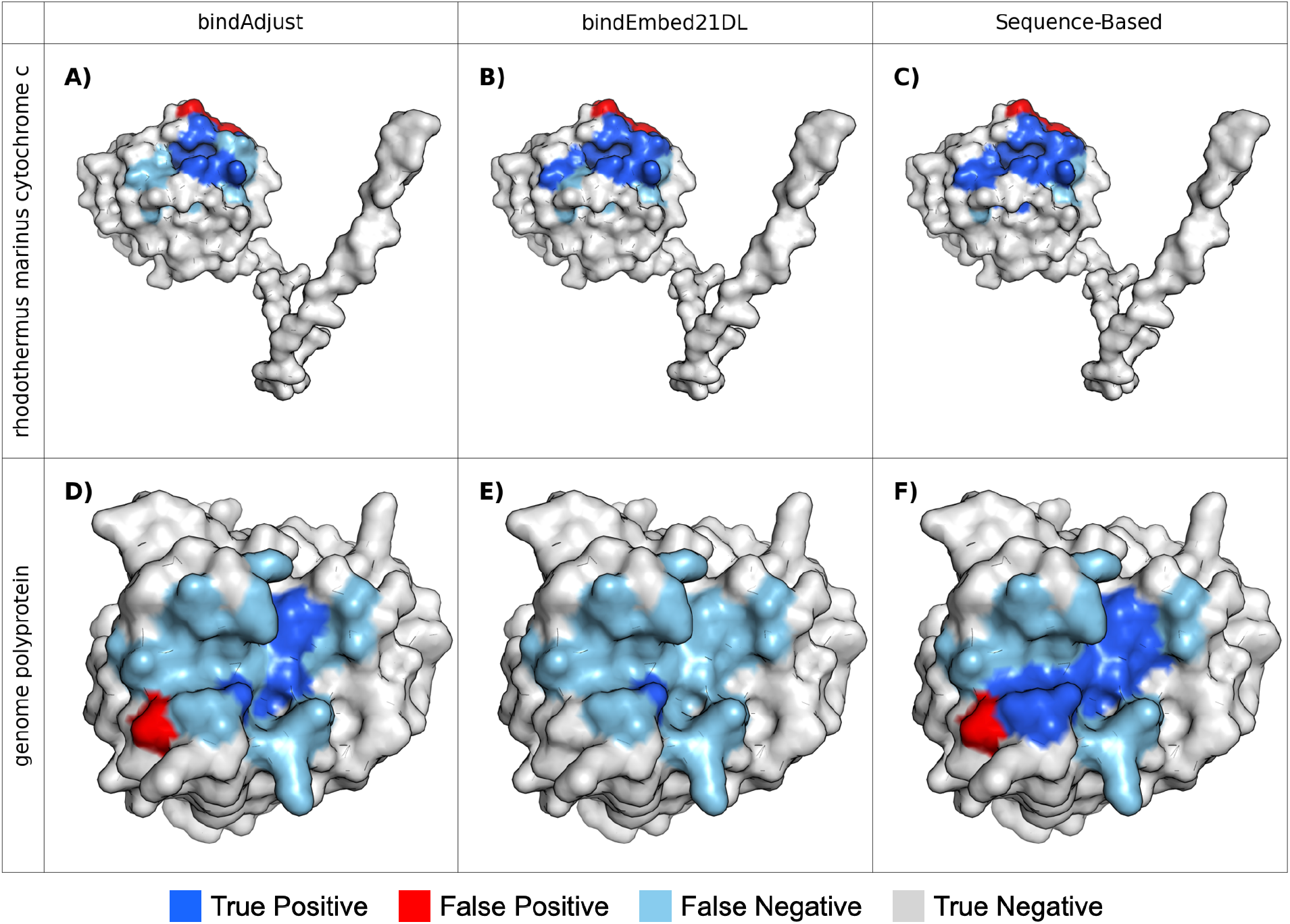
Visualization of two proteins comparing the prediction of bindEmbed21DL, bindAdjust and the sequence-sed refinement method. Rhodothermus marinus cytochrome c (UniProt ID B3FQS5 [27]), first row, and genome yprotein (UniProt ID E7E815 [28]), second row, are shown with predicted binding residues. <u>Dark blue:</u> Residues correctly dicted to bind to small ligand (True Positives), <u>Red</u>: Residues falsely predicted to bind (False Positives). <u>Light blue</u>: sidue false predicted to be non-binding (False Negatives), <u>Grey</u>: Residues correctly predicted to not bind (True gatives). For the rhodothermus marinus cytochrome c, the sequence-based method(B) slightly improves over dEmbed21DL(A) by adding five new binding predictions of which three are correct. bindAdjust (C=38) predicts seven re residues compared to bindEmbed21DL, resulting in 20 correct predictions out of 23 total predictions. For the genome yprotein, bindEmbed21DL(D) predicts six residues to bind. The sequence-based method(E) heavily underpredicts ding by (correctly) predicting only one residue to bind. bindAdjust(F) performs the best, it predicts 12 residues as binding which ten are correct.

For another protein, the genome polyprotein (Fig. 3, second row; UniProt ID E7E815 [28]), BioLiP annotates 21 residues as binding to a small molecule. bindEmbed21DL only predicts six residues as binding of which 5 are correct (Precision=83; Recall=24%; Fig. 3D), heavily underpredicting binding for this protein. On the other hand, bindAdjust predicts 12 residues as binding of which ten are correct (Precision=83%, Recall=48%; Fig. 3F). The sequence-based method is unable to improve upon the predictions of bindEmbed21DL but rather removes correct predictions of bindEmbed21DL Fig. 3 by predicting only one residue (correctly) as binding (Precision=100%, Recall=5%; Fig. 3E). The decrease in performance of the sequence-based method can be explained by the distance of the bindEmbed21DL predictions on the sequence (positions 39, 70, 146, 160, 163 and 167). bindAdjust, on the other hand, manages to extend the binding residue predictions of bindEmbed21DL by adding predictions close in structure to the six residues already identified by bindEmbed21DL resulting in a larger surface resembling an actual binding site (Fig. 3F).

## Conclusion

Here, we propose bindAdjust, a method to refine binding residue predictions using 3D structures. Structures are predicted using ColabFold [20] which builds upon AlphaFold2 [22]. bindAdjust incorporates the structural information in the form of a distance map into the predicted binding probabilities of bindEmbed21DL [16]. It achieves this by calculating structure bonus values for each residue by analyzing its distances to all other residues and their respective binding probabilities (Eqn. 1). These bonus values are then used to adjust the output probabilities of bindEmbed21DL.

Compared to the sequence-based method bindEmbed21DL [16], predicting residues binding to small molecules, metal ions, and nucleic acids, bindAdjust can improve the recall for small molecules while maintaining the same level of precision (Fig. 2, Table S4) by incorporating structural information. Predictions for individual proteins cannot be improved for binding to metal ions and nucleic acids. Here, bindAdjust rather identifies a subset of proteins where bindEmbed21DL achieves a particularly high recall (Table S6 & S7), probably focusing on proteins where the binding residues predicted by bindEmbed21DL are close in space and form a binding site.

Applying a refinement method similar to bindAdjust but solely using information about residues close in sequence not in 3D space cannot compete with the performance of bindAdjust for all three ligand classes (Fig. 1**Error! Reference source not found**., Tables S4, S6, and S7). This clearly demonstrates that bindAdjust leverages the 3D structure of the protein to enhance the quality of the prediction. Further investigation of two examples showcases that the binding predictions added by bindAdjust are in fact close to the predictions of bindEmbed21DL extending the single binding residues to form a larger surface or binding site (Fig. 3C & 2F). Since binding predictions close in 3D space are not necessarily close in sequence, the sequence-based refinement cannot improve the predictions for the two examples with sometimes even removing correct predictions (Fig. 3D). Fig. 3. The algorithm at the core of bindAdjust is straight forward and easily understood, only taking 5 seconds to calculate bindAdjust scores for 1010 proteins even on low performance CPUs. Without the requirement of time-consuming training or downloading of large models, the method is applicable to large corpora and enables proteome wide studies.

Due to its speed and simplicity, bindAdjust can be easily adapted and applied to any type of binding prediction. In general, any prediction task strongly tied to structure such as protein function, binding, or disorder could benefit from incorporating (predicted) structures through an approach like bindAdjust.

## Abbreviations used

3D: three-dimensional;
AFDB: AlphaFold Protein Structure Database;
FN: false negative(s);
FP: false positive(s);
MSA: Multiple Sequence Alignment;
NLP: Natural Language Processing;
pLM: protein Language Model;
TN: true negative(s);
TP: true positive(s);

## Declarations

### Ethics approval and consent to participate

Not applicable

### Consent for publication

Not applicable

### Availability of data and materials

All data and code are available via the public GitHub repository at https://github.com/Rostlab/bindadjust.

### Authors’ contributions

ML conceived the idea for the study; TO, KE, and ML devised the experimental setup; LE implemented the algorithm; LE and TO performed the data analyses. LE, TO, KE, and ML interpreted the results of the data analyses. KW contributed parts of the distance map pipeline. BR provided guidance and further ideas. LE and TO wrote the manuscript with the help of the other authors. All authors read and approved the final manuscript.

## Acknowledgements

Thanks to Tim Karl and Inga Weise (both TUM) for invaluable help with technical and administrative aspects of this work; to Michael Heinzinger for running ColabFold for the selected dataset. Also, thanks to the developers of AlphaFold2 (John Jumper and the team at DeepMind) and ColabFold (primarily Milo Mirdita, Konstantin Schütze and Martin Steinegger and his team) for making their methods publicly available and accessible to others. Last, but not least, thanks to all those who maintain public databases in particular Steven Burley (PDB, Rutgers), Ioannis Xenarios (Swiss-Prot, SIB, Geneva), and Yang Zhang (BioLiP, University of Michigan) and their crews.

