## Supplementary Material for "Refining Embedding-Based Binding Predictions by Leveraging AlphaFold2 Structures"

Supporting online material
for:

### Refining Embedding-Based Binding Predictions by Leveraging AlphaFold2 Structures

Leopold Endres, Tobias Olenyi, Kyra Erckert, Konstantin Weißenow, Burkhard Rost & Maria Littmann

### Table of Contents for Supporting Online Material

### Short description of Supporting Online Material

In this Supporting Online Material (SOM), we provide more details about the dataset (1.1) and the bindAdjust algorithm by providing a visualization (1.2). Table S2 provides an overview over used terms and variables to improve readability. Table S3 depicts an example of a constructed distance map generated from sequence.

In section 2.4, we show more results for the performance of bindAdjust, bindEmbed21DL, and the sequence-based refinement on different data sets. We extend the analysis of bindAdjust to two other ligand classes besides small molecules, namely metal ions, and nucleic acids (2.5). For all three ligand classes, the performance of bindAdjust was highly influenced by the choice of C (Fig. S2). Comparing the performance of bindAdjust and bindEmbed21DL on different sets (Tables S5 & S6) showed that bindAdjust rather identified a set of proteins with good predictions from bindEmbed21DL than improving predictions of individual proteins.

### 1 Material

#### 1.1 Dataset composition.

The data set was taken from bindEmbed21DL [1]. It consists of 1,314 proteins experimentally determined to bind to metal ions, nucleic acids, or small molecules. The binding annotations were extracted from BioLiP [2]. 1,014 proteins were used for training and hyperparameter optimization of bindEmbed21DL, 300 for testing. More details on the data set construction can be found in [1]. We removed four proteins (UniProt IDs: C8BD48, P84801, Q9NZV6, Q00277), as they contained unusual amino acids which were not included in the structure prediction leading to missing coordinates in the structure. This resulted in a final development set of 1,010 proteins and a test set of 300 proteins (Table S1).

**Table S1: Statistics on data set composition ***

| | **Development Set** | **Test Set** |
|---|---|---|
| **Total Size** | 1010 | 300 |
| **Small Molecules** | 603 | 220 |
| **Metal Ions** | 454 | 124 |
| **Nucleic Acids** | 108 | 66 |

* Counts for proteins annotated to bind to small molecules, metal ions, or nucleic acids (DNA/RNA) in the development and test set. Some proteins are annotated with multiple ligand classes.

##### 1.2 Visualization of bindAdjust structure bonus calculation.

Fig. S1 shows a visualization of the structure bonus calculation of bindAdjust (Eqn. 1 in the main text) for the prediction of small molecule binding of a dioxygenase (UniProt ID Q9REI7 [3]). The first, one-dimensional input are the binding probabilities from bindEmbed21DL for the small ligand class (Fig. S1a). The second input is the two-dimensional distance map of the protein predicted by ColabFold [4] (Fig. S1b). The distance map records the distances between the C-alpha atoms of all residues of the protein in Angstrom and was derived from the predicted 3D conformation of the protein. Dividing the probability vector (Fig. S1a) by the residue distances (Fig. S1b) yields a matrix which contains structure influence values for each residue pair (Fig. S1c). The structure bonus (Fig. S1d) for this protein and specific ligand is the mean across the columns of the matrix (Fig. S1c).

The bonus values are added to the binding probabilities of bindEmbed21DL after being multiplied with a variable coefficient C. This results in the final bindAdjust scores (Eqn. 2 in the main text) which encode information from both the binding probabilities and the predicted structure of the protein.

**Fig. S1: Example calculation of bindAdjust structure values. ***

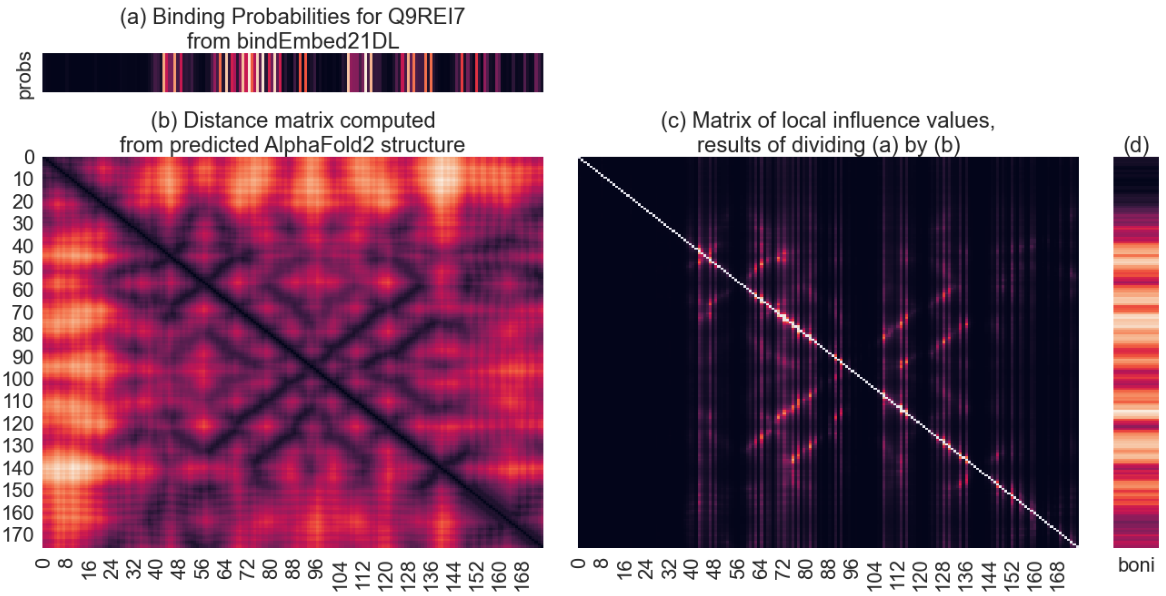

* (a) small binding probabilities for 2,4'-dihydroxyacetophenone dioxygenase (UniProt ID Q9REI7 [3]) of bindEmbed21DL [1], (b) distance matrix between C-alpha atoms computed from predicted ColabFold [4] structure, (c) matrix of local influence values resulting from the division of binding probabilities by the distance matrix, (d) final bonus values of bindAdjust, mean of matrix (c) across columns. Bonus values are added to original probabilities with a coefficient C. This way, predicted structural information is used to enrich the binding probabilities, resulting in a higher overall performance.

##### 1.3 Explanation of variables and scores.

**Table S2: Definition of variables, scores, and terms. ***

| | |
|---|---|
| bindEmbed21DL binding probabilities | Output of bindEmbed21DL, per residue, per ligand class binding probabilities |
| bindAdjust scores | Continuous, unnormalized output of bindAdjust, results of combining bindEmbed21DL with calculated structure bonus |
| Coefficient C | Coefficient applied to bindAdjust bonus before adding them to bindEmbed21DL binding probabilities |
| Cutoff | Threshold to classify whether a residue is considered binding or non-binding. If probability/score<=cutoff, the residue is considered non-binding; otherwise, it is considered binding. |

* We list and explain scores and variables commonly used throughout this work.

##### 1.4 Sequence-based refinement.

**Table S3: Example distance map constructed from sequence ***

| | **$R_1$** | **$R_2$** | **$R_3$** | **$R_4$** | **$R_5$** |
|---|---|---|---|---|---|
| **$R_1$** | 0 | 1 | 2 | 3 | 4 |
| **$R_2$** | 1 | 0 | 1 | 2 | 3 |
| **$R_3$** | 2 | 1 | 0 | 1 | 2 |
| **$R_4$** | 3 | 2 | 1 | 0 | 1 |
| **$R_5$** | 4 | 3 | 2 | 1 | 0 |

* Generated sequence-based distance map for a protein containing five residues. The distance in between neighbouring residues was set to 1.

### 2 Additional Results

#### 2.1 Performance of bindEmbed21DL and bindAdjust on development set. *

**Fig. S2: Average performance metrics of bindEmbed21DL, the sequence-based refinement, and bindAdjust for development set and parameter settings for the prediction of small molecule binding. ***

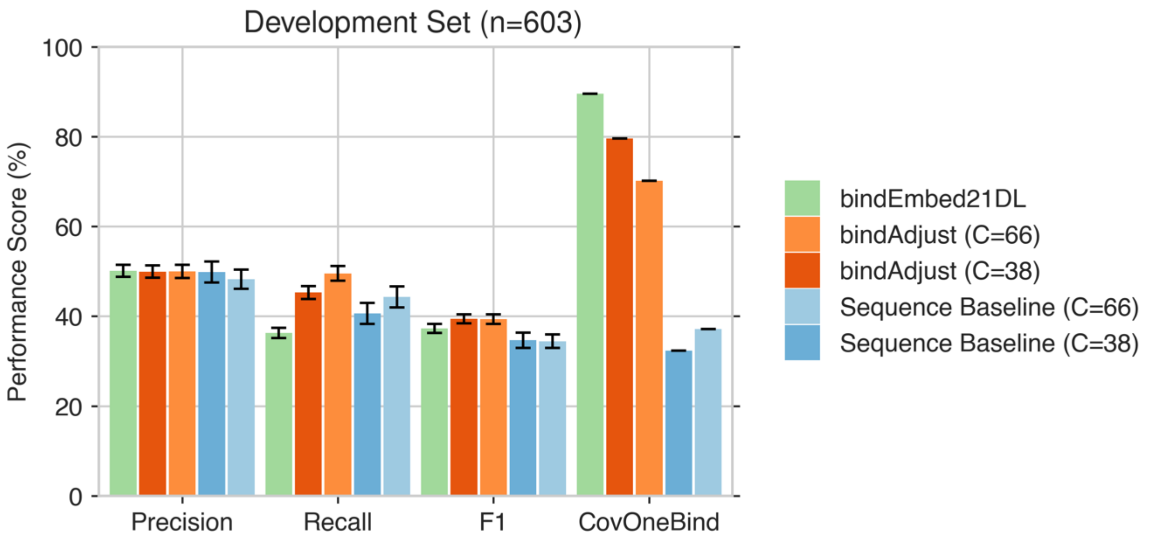

* *DevSet603*: 603 proteins binding to small molecules from the development set of bindEmbed21DL. For the chosen values for C (38, 66), only for C=38 the sequence-based refinement can reach the target precision of 50%. bindAdjust achieves a recall of 45±1% and 50±2, respectively, significantly improving over bindEmbed21DL.

#### 2.2 Optimization of C for binding to small molecules, metal ions, and nucleic acids.

**Fig. S3: Performance of bindAdjust for different values of C. ***

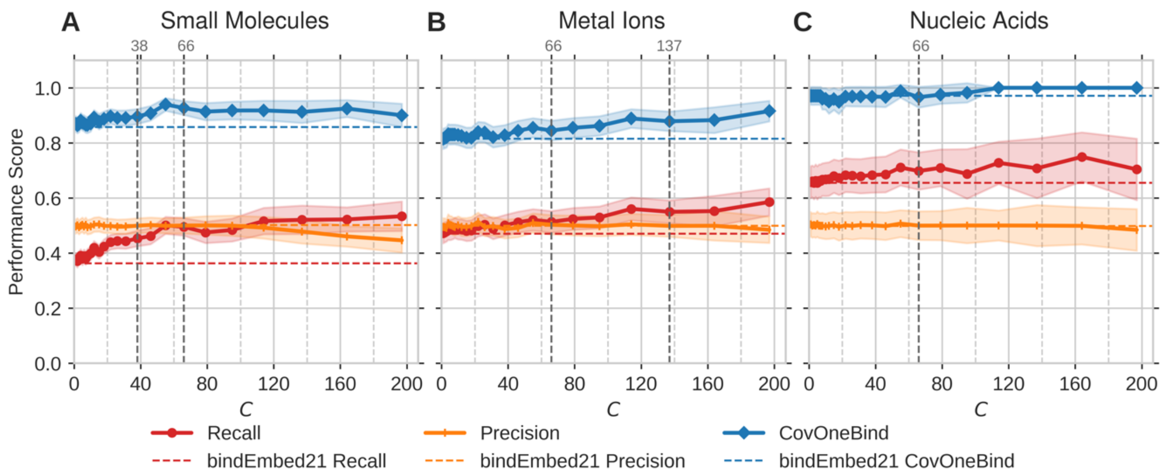

* Precision, recall, and CovOneBind of bindAdjust on the development set. The model was evaluated using values for C ranging from 0 to 197 with increasing intervals. We evaluated the performance of bindAdjust for the three ligand classes separately: **A.** small molecules, **B.** metal ions, and **C.** nucleic acids. Dashed horizontal lines indicate the performance of bindEmbed21DL. Dashed vertical lines mark the C values which we use for further analyses. For each parameter selection, the cutoff to classify a residue as binding/non-binding was adjusted to obtain an average precision of 50%. At this cutoff, CovOneBind and mean recall were recorded. For C>70, C>150, and C>137 bindAdjust cannot achieve the target precision of 50% for small molecules, metal ions, and nucleic acids, respectively.

##### 2.3 Performance of bindEmbed21DL and bindAdjust on subsets of development set. *

**Fig. S4: Average performance metrics of bindEmbed21DL and bindAdjust(C=38) for the prediction of small molecule binding on protein subsets identified by bindAdjust. ***

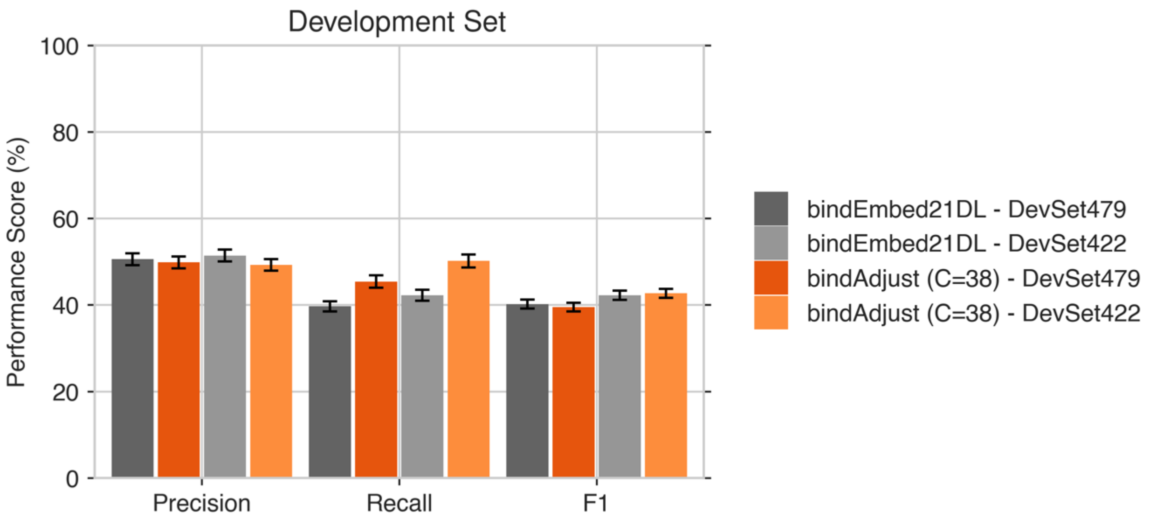

* DevSet479: 479 proteins of the development set binding to small molecules for which both bindEmbed21DL and bindAdjust(C=38) predict at least one residue as small binding. DevSe422: 422 proteins of the development set binding to small molecules for which bindAdjust(C=66) predicts at least one residue as small binding. Using DevSet422, both bindAdjust(C=38) and bindEmbed21DL perform better than for DevSet479. This demonstrates that bindAdjust and large values for C can be used to identify a subset of proteins for which both analysed models predict small binding exceptionally well.

#### 2.4 Performance evaluation of bindEmbed21DL, bindAdjust, and the sequence-based refinement on different data sets for small molecule binding.

**Table S4: Performance comparison for small ligand binding of bindEmbed21DL, the sequence-based refinement, and bindAdjust on development and test set. ***

| Dataset | Method | C | Cutoff | Precision | Recall | F1 Score | CovOneBind |
|---|---|---|---|---|---|---|---|
| DevSet603 | bindEmbed21DL | | 0.65 | 50±1% | 36±1% | 37±1% | 90% |
| | bindAdjust | 38 | 1.15 | 50±1% | 45±1% | 39±1% | 80% |
| | | 66 | 1.55 | 50±1% | 50±2% | 39±1% | 70% |
| | Sequence-based refinement | 38 | 1.35 | 50±1% | 40±2% | 35±2% | 70% |
| | | 66 | 1.95 | 45±2% | 43±2% | 33±1% | 57% |
| TestSet220 | bindEmbed21DL | | 0.55 | 50±2% | 47±2% | 44±2% | 94% |
| | bindAdjust | 38 | 0.95 | 50±2% | 52±2% | 44±2% | 91% |
| | | 66 | 1.30 | 50±2% | 53±2% | 44±2% | 82% |
| | Sequence-based refinement | 38 | 1.05 | 50±2% | 45±2% | 40±1% | 88% |
| | | 66 | 1.45 | 50±2% | 45±2% | 38±2% | 78% |

* Average performance metrics of bindEmbed21DL, the sequence-based refinement, and bindAdjust for different sets and parameter settings for the prediction of small molecule binding. DevSet603: 603 proteins binding to small molecules from the development set of bindEmbed21DL. TestSet220: 220 proteins binding to small molecules from the test set. For both sets, cutoffs were chosen so that the methods achieved around 50% precision leading to different cutoffs for the development and the test set. Standard errors are given as error estimates.

**Table S5: Performance comparison for small ligand binding of bindEmbed21DL and bindAdjust on subsets of the development and test set identified by bindAdjust. ***

| | Dataset | Cutoff | Precision | Recall | F1 Score | CovOneBind |
|---|---|---|---|---|---|---|
| **bindEmbed21DL** | DevSet479 | 0.65 | 51±1% | 40±1% | 40±1% | 100% |
| **bindAdjust (C=38)** | DevSet479 | 1.15 | 50±1% | 45±1% | 39±1% | 100% |
| **bindEmbed21DL** | DevSet422 | 0.65 | 51±1% | 42±1% | 42±1% | 100% |
| **bindAdjust (C=38)** | DevSet422 | 1.15 | 49±1% | 50±2% | 43±1% | 100% |
| **bindEmbed21DL** | TestSet199 | 0.55 | 51±2% | 48±2% | 45±1% | 100% |
| **bindAdjust (C=38)** | TestSet199 | 0.95 | 50±2% | 52±2% | 45±2% | 100% |
| **bindEmbed21DL** | TestSet180 | 0.55 | 51±2% | 50±2% | 47±2% | 100% |
| **bindAdjust (C=38)** | TestSet180 | 0.95 | 48±2% | 56±2% | 47±1% | 100% |

* Models benchmarked on the 479 small binding proteins of the development set (DevSet479) for which both models predict at least one residue to bind. A second benchmarking set (DevSet422) was obtained by applying bindAdjust (C=66) and using the proteins for which this model makes predictions. The same procedure was applied to the test set. TestSet199 are the 199 proteins for which bindAdjust (C=38) predicts small binding. TestSet180 is another subset of proteins identified by increasing the C parameter to 66. Residues with a bindEmbed21DL output probability/bindAdjust score > cutoff were classified as binding. Standard errors are given as error estimates.

##### 2.5 bindAdjust identifies subset of proteins with good predictions for metal ions and nucleic acids.

**Performance for metal ion binding.**

454 of the 1010 proteins in the development set have at least one residue annotated as metal binding by BioLiP [2]. For 418 of those (CovOneBind=92%), bindEmbed21DL achieves recall=47±2% for a precision of 51±2%. While the increase in performance is not as pronounced as for the small ligand class (see main text), bindAdjust manages to improve that recall. As for the small class, higher values for C lead to an increase in recall and a decrease in CovOneBind (Fig. S3b) while for C>150, bindAdjust cannot achieve the target precision anymore. Also, for C<31, we do not observe any improvement. C=66 and C=137 are selected for further analysis and the recall is recorded at an average precision of 50%. C=66 increases the recall to 51±2% at CovOneBind of 83%; for C=137, it rises even further to 55±2% at CovOneBind of 72% (Fig. S3b). On the other hand, the sequence-based method

yields recall=48±2% (Table S6) for C=66 and cannot reach the target precision of 50% for C=137 (Fig. S3b), i.e., the sequence-based method cannot improve over bindEmbed21DL while bindAdjust at C=66 leads to a slight improvement.

For a better comparison, we evaluate the performance of bindEmbed21DL and bindAdjust for C=66 on three different datasets: (i) all metal-binding proteins, (ii) the 379 proteins for which bindAdjust (C=66) and bindEmbed21DL predicted at least one residue as metal binding, and (iii) the 329 proteins for which bindAdjust (C=137) predicted at least one residue as metal binding (Table S6). We use this last dataset to check whether we can use larger values for C to identify a subset of proteins for which both bindEmbed21DL and bindAdjust (C=66) perform well. In fact, both bindAdjust (C=66) and bindEmbed21DL achieve a higher performance on $DevSet_{C=137}$ than on $DevSet_{C=66}$ with bindAdjust achieving a slightly higher recall than bindEmbed21DL (Table S6) indicating that bindAdjust is rather able to zoom into the subset of proteins with good predictions from bindEmbed21DL than improving predictions for individual proteins.

**Table S6: Average performance for metal binding protein in the development set for bindAdjust using C=66 and C=137. ***

| Set | Method | Cutoff | Precision | Recall | F1 | CovOne Bind |
|---|---|---|---|---|---|---|
| **DevSet454** | **bindEmbed21DL** | 0.375 | 51±2% | 47±2% | 42±2% | 93% |
| | **bindAdjust (C=66)** | 0.775 | 51±2% | 51±2% | 40±2% | 85% |
| | **Sequence-Based Method (C=66)** | 1.125 | 50±2% | 48±2% | 38±2% | 67% |
| **$DevSet379_{C=66}$** | **bindEmbed21DL** | 0.375 | 54±2% | 51±2% | 45±2% | 100% |
| | **bindAdjust (C=66)** | 0.775 | 51±2% | 51±2% | 40±2% | 100% |
| **$DevSet329_{C=137}$** | **bindEmbed21DL** | 0.375 | 55±2% | 55±2% | 49±2% | 100% |
| | **bindAdjust (C=66)** | 0.775 | 51±2% | 57±2% | 44±2% | 100% |

* Performance comparison in terms of precision, recall, and F1 score between bindAdjust (C=66) and bindEmbed21DL using different protein sets for evaluation. *DevSet454*: 454 proteins binding to metal ions from the development set. *$DevSet379_{C=66}$*: 379 proteins for which bindAdjust with C=66 and bindEmbed21DL predict at least one residue as metal binding. *$DevSet329_{C=137}$*: 329 proteins for which bindAdjust with C=137 and bindEmbed21DL predict at least one residue as metal binding. For DevSet454, we use the cutoffs resulting in an average precision of 50%. We use the same respective cutoffs on the other two sets. Standard errors obtained from bootstrapping are given as error estimates.

**Performance for nucleic acid binding.**

108 of the 1010 proteins of the development set have at least one residue annotated to bind to nucleic acids (DNA or RNA) in BioLiP [2]. For 104 of those proteins, bindEmbed21DL makes a prediction (CovOneBind=94%) achieving recall=64±3% at a precision of about 50% (Fig. S3c). As for the two other ligand classes, the parameter C exerts a substantial influence of the methods performance (Fig. S3c). bindAdjust (C=66) increases the recall of the predictions by four percentage point to 70±4% (Table S7). The sequence-based refinement is only able to increase the recall to 63±4% (Table S7). We also compare the performance of both methods using only the 88 proteins which are predicted to bind to nucleic acids by both (DevSet88). For these proteins, bindEmbed21DL achieves a recall of 73±2% outperforming bindAdjust (recall=70±4%; Table S7). This shows that, similar to the result for metal ion binding, bindAdjust cannot boost the overall performance of the binding predictions of bindEmbed21DL for nucleic acids, but it can be used to identify a subset of proteins with particularly good binding predictions from bindEmbed21DL.

**Table S7: Average performance for nucleic acid binding proteins in the development set for bindAdjust using C=66. ***

| Set | Method | Cutoff | Precision | Recall | F1 | CovOne Bind |
|---|---|---|---|---|---|---|
| DevSet108 | bindEmbed21DL | 0.375 | 50±2% | 66±3% | 51±2% | 96% |
| | bindAdjust (C=66) | 1.350 | 50±3% | 70±4% | 50±2% | 81% |
| | Sequence-Based Method (C=66) | 2.250 | 50±3% | 63±4% | 48±3% | 64% |
| DevSet88$_{C=66}$ | bindEmbed21DL | 0.375 | 49±2% | 73±2% | 56±2% | 100% |
| | bindAdjust (C=66) | 1.350 | 50±3% | 70±4% | 50±2% | 100% |

* Performance comparison in terms of precision, recall, and F1 score between bindAdjust (C=66) and bindEmbed21DL using different protein sets for evaluation. *DevSet108:* 108 proteins binding to nucleic acids from the development set. *DevSet88$_{C=66}$*: 88 proteins for which bindAdjust with C=66 and bindEmbed21DL predicted at least one residue as nucleic acid binding. Standard errors are given as error estimates.
